# Quantitative and temporal measurement of dynamic autophagy rates

**DOI:** 10.1101/2021.12.06.471515

**Authors:** Nitin Sai Beesabathuni, Priya S. Shah

**Affiliations:** Department of Chemical Engineering, University of California, Davis; Department of Microbiology and Molecular Genetics, University of California, Davis

**Keywords:** Autophagy flux, bafilomycin A1, cargo, live cell fluorescence microscopy, non-steady state, puncta, rapamycin, steady state, temporal dynamics, wortmannin

## Abstract

Autophagy is a multistep degradative process that is essential for maintaining cellular homeostasis. Systematically quantifying flux through this pathway is critical for gaining fundamental insights and effectively modulating this process that is dysregulated during many diseases. Established methods to quantify flux use steady state measurements, which provide limited information about the perturbation and the cellular response. We present a theoretical and experimental framework to measure autophagic steps in the form of rates under non-steady state conditions. We use this approach to measure temporal responses to rapamycin and wortmannin treatments, two commonly used autophagy modulators. We quantified changes in autophagy rates in as little as 10 minutes, which can establish direct mechanisms for autophagy perturbation before feedback begins. We identified concentration-dependent effects of rapamycin on the initial and temporal progression of autophagy rates. We also found variable recovery time from wortmannin’s inhibition of autophagy, which is further accelerated by rapamycin. In summary, this new approach enables the quantification of autophagy flux with high sensitivity and temporal resolution and facilitates a comprehensive understanding of this process.

## Introduction

Macroautophagy (hereafter referred as autophagy) is an intracellular recycling process that breaks down misfolded proteins and damaged organelles into their primary building blocks. This dynamic process involves autophagosome formation, the fusion of autophagosomes with lysosomes, and the turnover of autolysosomes. Constitutive autophagy is important for cellular homeostasis and is modulated during many extrinsic stresses such as nutrient deprivation or pathogen infection [1,2]. This process is also dysregulated during chronic diseases associated with aging, neurodegeneration and cancer [3,4]. Autophagy can be modulated using pharmacological agents and is a major drug development target for treating cancer, neurodegeneration, and pathogen infection [5,6]. Along with medical applications, autophagy modulation has also shown potential to enhance biomanufacturing by increasing cell longevity [7]. Thus, quantifying autophagy dynamics will be critical in any application that involves modulating this process.

A major challenge in determining how autophagy is modulated is the limited tools that allow for direct and absolute quantification of each step in the process. Western blot is commonly used to estimate autophagy; however, it is typically less quantitative and less sensitive. Western blot also does not allow measurement of all the autophagic steps. Fluorescent reporter systems have enabled the quantification of autophagosomes in live cells. Green fluorescent protein (GFP) labeling of MAP1LC3/LC3, a protein associated with autophagosomes and autolysosomes, allows for quantification of autophagosome accumulation. GFP is pH sensitive and is bleached in the acidic environment of the autolysosome in this system [8,9]. Tandem green and red (GFP:RFP) labeling of LC3 has also be used to identify acidic autolysosomes since RFPs are acid-stable [10,11]. Even though these systems allow direct monitoring of autophagosomes and autolysosomes in real-time, they do not provide direct quantification of each step, also known as ‘autophagy flux’. For instance, the accumulation of autophagosomes could be a result of an increase in the formation or decrease in the clearance of autophagosomes. The same principle applies to autolysosomes.

To quantitatively measure autophagy flux, the inputs and outputs of autophagosome and autolysosome accumulation must be dissected. Measuring autophagosome accumulation after inhibiting clearance with small molecules such as bafilomycin A1 is a commonly used approach for measuring autophagic flux [12–14]. Nevertheless, this approach does not provide a direct quantification of autophagosome and autolysosome clearance steps. A new approach using a novel fluorescent probe was described to quantify autophagic flux by measuring the GFP:RFP signal ratio without adding lysosomal inhibitors [15]. However, this method does not provide a direct quantification of autolysosomes. It is also not sensitive to identifying differences in the autophagic flux if the changes are relatively similar. For example, inhibition of autophagosome formation could lead to similar changes in both GFP and RFP signals. Thus, a true change in flux may still result in a similar GFP:RFP ratio as basal, leading to ambiguity. Another case could be equal and simultaneous initiation and inhibition of autophagosome formation and clearance (resulting in constant GFP signal), which could lead to no observable difference in the fluorescence levels. Finally, most measurements made using the methods discussed above are made long after perturbation, when the system reaches a new steady state where the rates of all the steps are equal. Although this provides very useful information about the final autophagy state, it is incapable of informing the nature of the perturbation and the dynamic response of the cells to the perturbation. To illustrate, an autophagosome formation inhibitor and an autophagosome formation inducer might reach the same final steady state even though they perturb autophagy very differently. Therefore, it is essential to temporally quantify all the steps involved in autophagy to gain a better understanding about the perturbation as well as the regulatory mechanisms of this dynamic process.

Here, we present an approach to quantify the rates of each step of the autophagy pathway under non-steady state conditions. This approach directly builds upon previous advances that provided a quantitative framework to measure autophagy rates at a steady state [12,16], and is enabled by high-throughput live cell imaging and measurement of instantaneous autophagy rates. With this approach, we study the effects of well-characterized autophagy regulators rapamycin and wortmannin. Through our non-steady state analysis, we show that rapamycin dynamically regulated autophagy flux to reach an elevated state that decreases back to basal levels over time. We use the non-steady state rate approach to reveal two modes of regulation that cause this dynamic behavior. We further show that rapamycin concentrations can be used to precisely modulate autophagy flux. On the other hand, we show wortmannin initially inhibits autophagy flux,which is recovered over time in a concentration-dependent manner. Taken together, this innovative approach has the potential to provide novel insights related to autophagy and mechanisms driving autophagy-regulating perturbations through quantitative measurements.

## Results

### Non-steady state measurement of autophagy rates

Measurement of steady state autophagy flux has long been performed [17,18]. Loos and colleagues established a formal framework to quantify autophagy flux or the rate of autophagosome formation under steady state conditions [12,16]. The model considers the whole autophagic process as a multistep process governed by three steps: 1) the rate of formation of autophagosomes (R_1_), 2) the rate of autolysosome formation via fusion of autophagosomes with lysosomes (R_2_) and 3) the rate of degradation of autolysosomes (R_3_) (**Fig 1A**). This method relies on quantifying autophagosomes and their accumulation over time in live cells by using GFP-labeled LC3 as described above. Performing a mass balance on the autophagosomes (AP) yields an expression for the rate of change of autophagosomes:

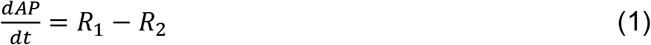

**Figure 1.**
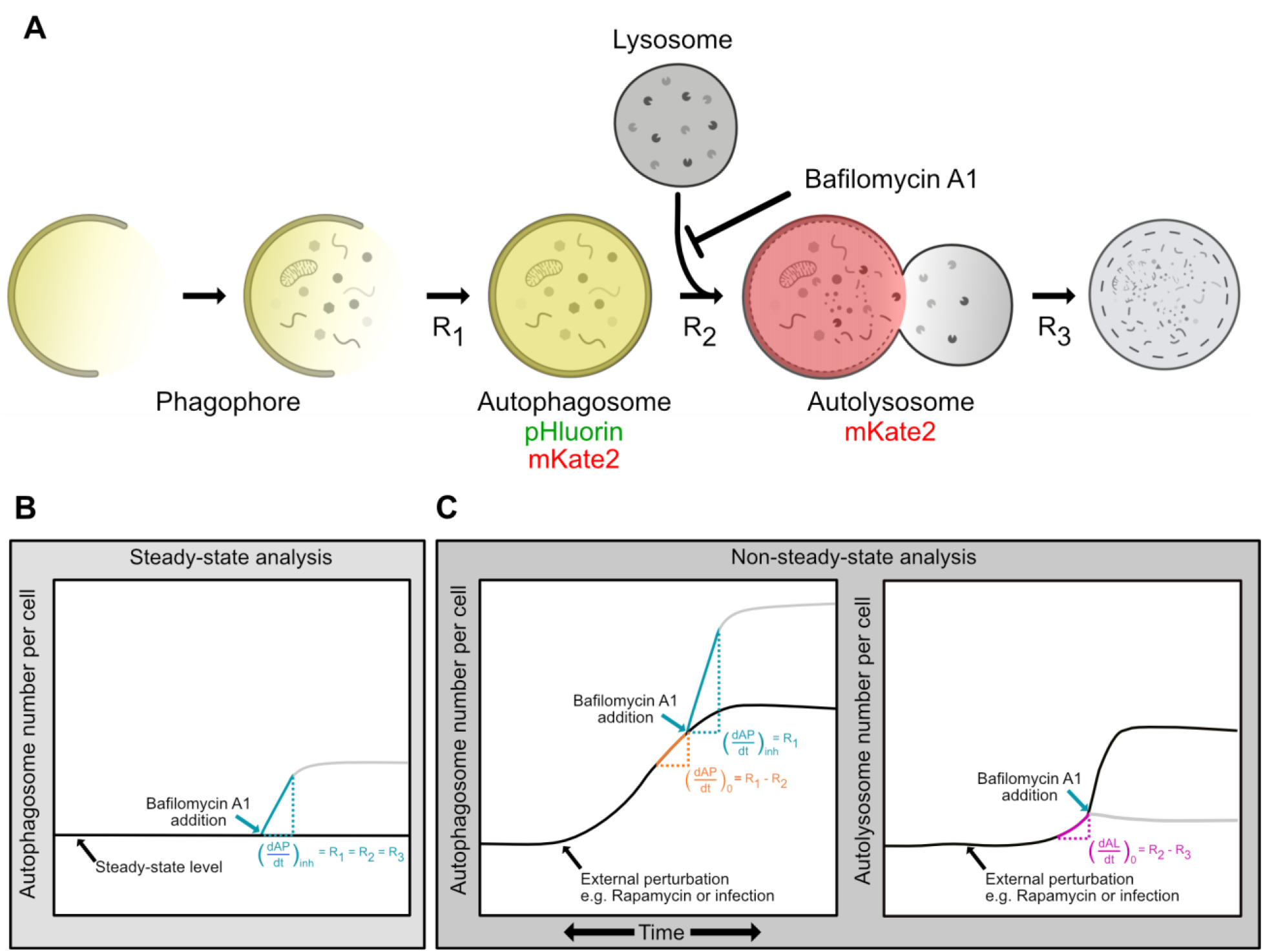
Conceptual framework of non-steady state analysis of autophagy rates. **(A)** Phagophores elongate to form autophagosomes. Autophagosomes fuse with lysosomes to form autolysosomes. Contents are degraded in autolysosomes. The rates of each of these steps (R_1_, R_2_, and R_3_) can be measured using a mass action model and live-cell imaging. Fluorescently tagged LC3 (pHluorin-mKate2-LC3) can be used to quantify autophagosomes (pHluorin- and mKate2-positive) and autolysosomes (mKate2-positive, pHluorin is quenched at low pH). **(B)** Measurement of autophagosome numbers following inhibition of autophagosome-lysosome fusion using bafilomycin A1 allows for measurement of R_1_, the rate of autophagosome formation. When performed at a steady state, this rate is equal to the other rates in the pathway. **(C)** When changes in autophagosome and autolysosome numbers are measured using an instantaneous rate approach, all rates in the autophagy pathway (R_1_, R_2_, and R_3_), which may not be equal under dynamic conditions, can be measured.

Similarly, the rate of change of autolysosomes (AL) can be written as:

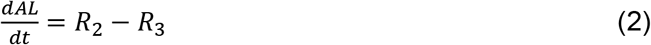

Under steady state, no change in autophagosomes or autolysosomes over time is observed because R_1_, R_2_, and R_3_ are equal. Using bafilomycin A1 to inhibit the fusion of the autophagosome with lysosomes sets *R*_2_ = 0 and results in autophagosome accumulation. Immediately post inhibition:

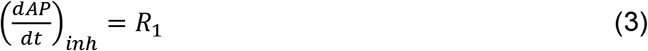

where 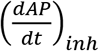 is the accumulation rate of autophagosomes following inhibition of R_2_ by bafilomycin A1 [12,16] (**Fig 1B**).

Despite the success of quantitative autophagy flux measurements, non-steady state measurements (dynamic conditions in which R_1_, R_2_, and R_3_ may not be equal) remain out of reach. We expanded on this previous work to develop a non-steady state rate approach that enables the evaluation of all three rates under dynamic conditions (**Fig 1C**). Measuring the change in the number of autophagosomes with time just before the chemical inhibition would provide the net rate of change of autophagosomes at that time point 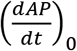. R_2_ can then be evaluated using equations (1) and (3):

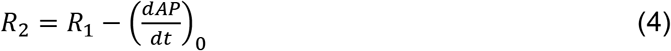

We can extend this analysis to also quantify the instantaneous net rate of change of autolysosomes 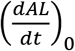 prior to chemical inhibition. R_3_ can be evaluated using equations (2) and (4):

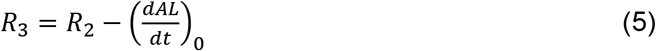

Thus, we can recover the absolute formation and fusion rates of autophagosomes along with the degradation rate of autolysosomes as a function of time by carrying out this approach at multiple time points.

### Experimental system to monitor autophagy dynamics

Accurately quantifying autophagy rates over time requires a method to distinguish autophagosomes from autolysosomes and to track them in live cells simultaneously. Several tandem reporter systems that can distinguish autophagosomes from autolysosomes have been previously described [19–21], though none have been used to extract rate data for all steps in the autophagy pathway. We used the previously developed Super-Ecliptic, pHluorin-mKate2-LC3 system [11]. pHluorin is an acid-sensitive GFP, while mKate2 is an acid-stable RFP. Thus, autophagosomes are green and red, while the acidic autolysosomes are only red (**Fig 1A**).

We first confirmed that accumulation of pHluorin-mKate2-labeled puncta is specific to autophagosomes using the tandem reporter with wild-type (WT) LC3 and a LC3 mutant (LC3 ΔG). LC3 ΔG lacks the glycine at the carboxyl-terminus, which is essential for proper lipidation and association with autophagosomes [8,22]. The addition of bafilomycin A1 and rapamycin, two well-established modulators of autophagy [23,24], lead to the expected accumulation of pHluorin-mKate2-labeled puncta in cells expressing WT LC3, but not in LC3 ΔG-expressing cells (**Fig 2A**).

**Figure 2.**
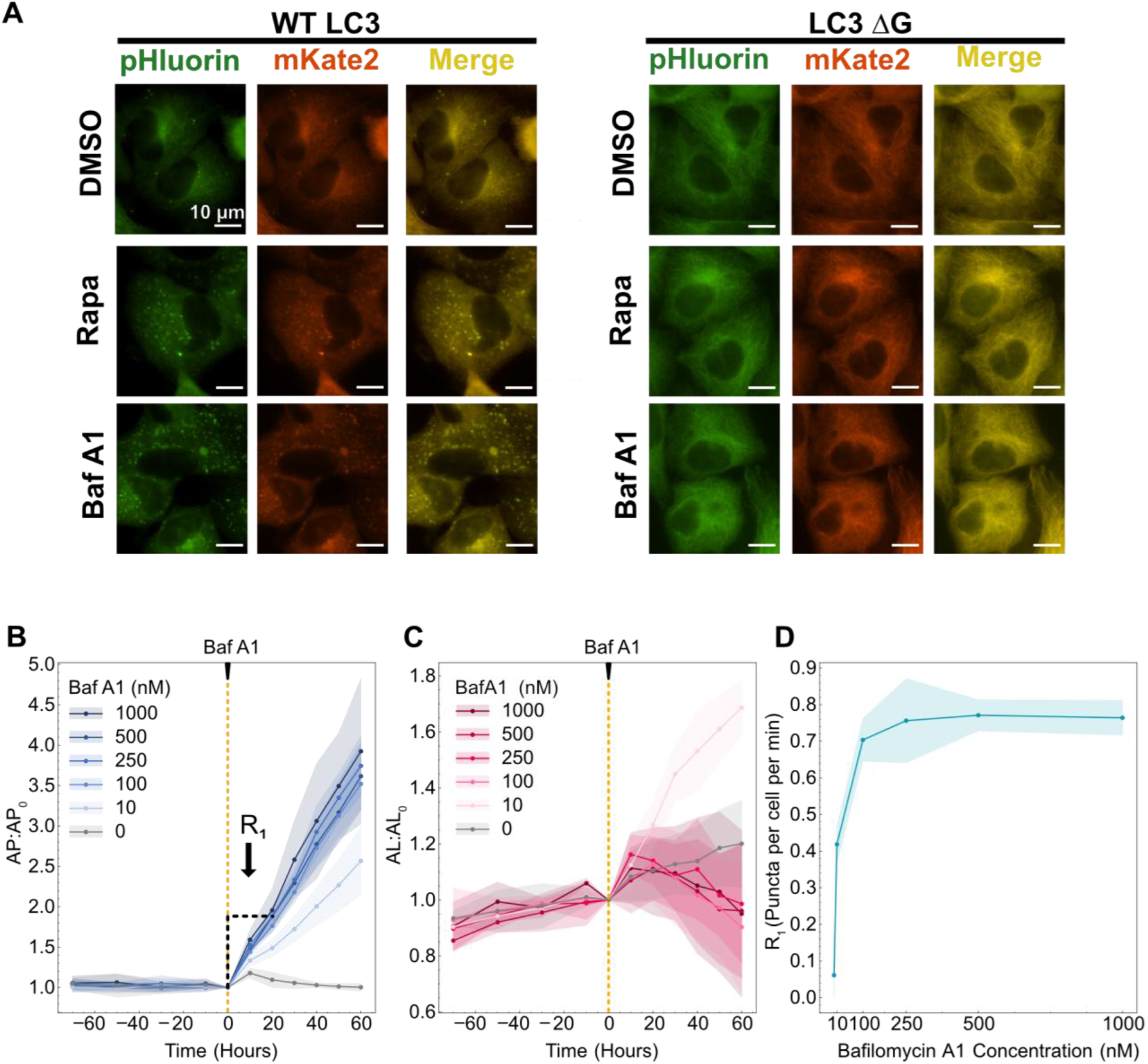
Calibration of system conditions for data collection. **(A)** Images of cells expressing the pHluorin-mKate2-LC3 tandem fluorescent reporter following DMSO, 100 nM rapamycin (Rapa) and 500 nM bafilomycin A1 (Baf A1) treatment. WT LC3 represents wild type and LC3 ΔG was used as a negative control since this mutant cannot be lipidated for phagophore association. **(B)** Autophagosomes and **(C)** autolysosomes were quantified over 90 minutes before the addition of bafilomycin A1 and 60 minutes after. R_1_ was calculated using the first 20 minutes of data following bafilomycin A1 treatment. **(D)** R_1_ rates are plotted as a function of bafilomycin A1 concentration.

We then calibrated our experimental system to determine the optimal concentration of bafilomycin A1 to inhibit R_2_ completely. We monitored autophagosome and autolysosome dynamics over time before and after the addition of bafilomycin A1 at various concentrations (**Fig 2B and C**). Autophagosome and autolysosome numbers were confirmed to be at a steady state prior to the addition of bafilomycin A1. We observed a constant increase in autophagosomes over time following the addition of bafilomycin A1 for all concentrations tested, with more dramatic increases at higher concentrations. For autolysosomes, we did not observe any considerable changes for higher concentrations (100 nM and above) but for 10 nM bafilomycin A1 we observed an increase in the autolysosome numbers. This could be due to complete inhibition of autolysosome clearance by bafilomycin A1 [17,23] but only partial inhibition of the fusion step as indicated by the lower slope of autophagosome increase, leading to continuous autolysosome production but no clearance.

We used autophagosome data to determine R_1_ as a function of bafilomycin A1 concentration. R_1_ was measured as the slope using 20 minutes of data immediately following the addition of bafilomycin A1 (**Fig 2B**). The ability to measure rates within 20 minutes is a major advantage of this system as it allows the measurement of instantaneous R_1_ with minimal feedback from bafilomycin A1 addition. A saturation of R_1_ was observed starting at 100 nM bafilomycin A1 (**Fig 2D**). However, to ensure complete inhibition of R_2_ even during induced conditions (*e.g*., higher autophagosome and lysosome numbers), a concentration of 500 nM bafilomycin A1 was used for all subsequent experiments.

### Rapamycin-induced autophagosome and autolysosome dynamics are concentration-dependent

We next tested the ability to monitor autophagosome and autolysosome temporal dynamics using rapamycin, which induces autophagosome formation through the inhibition of mTORC1 [25–27]. We tested seven concentrations of rapamycin (**Fig 3, S1A and B**). Autophagosomes increased following rapamycin treatment, with higher rapamycin concentrations resulting in a more rapid increase (**Fig 3A, S1A**). For the highest concentrations of rapamycin (≥ 10 nM), this rapid increase peaked at 30 minutes post-treatment, followed by a gradual decrease. We observed a saturation behavior for concentrations above 10 nM. A mid-range concentration of rapamycin (1 nM) resulted in a more gradual increase in autophagosome numbers, followed by a slight decrease. The lowest concentration of rapamycin tested (0.1 nM) had no effect. Autolysosome dynamics followed similar concentration-dependent trends, with slightly delayed peaks at 1.5-2 hours post-treatment for higher concentrations (**Fig 3B, S1B**). Interestingly, there was no difference in autophagosome and autolysosome peak time (3.5-4 hours) for the mid-range concentrations of rapamycin (0.5 and 1 nM). The raw autophagosome and autolysosome data are provided in the supplemental (**Fig S1C and D**). We also confirmed rapamycin inhibition of mTORC1 activity by monitoring change in the phosphorylation status of S6 ribosomal protein at the Ser 240/244 site, which is a downstream substrate of mTORC1 (**Fig S1E**).

**Figure 3.**
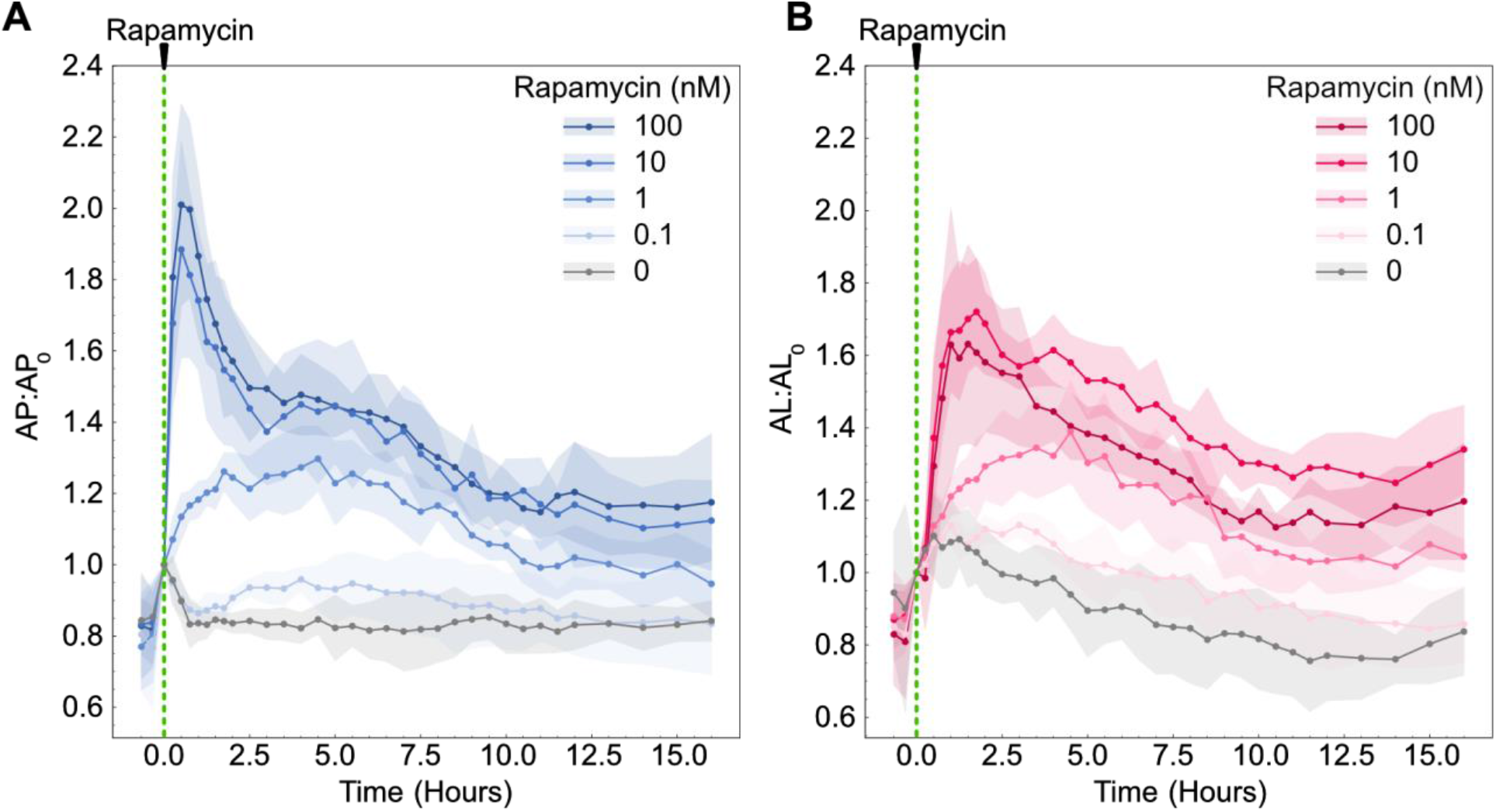
Autophagosome and autolysosome dynamics are a function of rapamycin concentration. **(A)** Autophagosome and **(B)** autolysosome number dynamics after rapamycin treatment. The indicated concentration of rapamycin was added at 0 minutes. The number of autophagosomes and autolysosomes at 0 minutes was used as the normalization factor. Data points represent mean while shaded area represents ± standard deviation. Four independent replicates were performed.

### Time evolution of autophagy rates reveals initial rate-limiting steps

The autophagosome dynamics for high concentrations of rapamycin (≥ 10 nM) suggested two possible models of cellular response to rapamycin treatment. The sudden increase in autophagosome numbers followed by a decrease could be driven by a rapid increase in the rate of autophagosome formation (R_1_), while the rates of autolysosome formation and degradation (R_2_ and R_3_, respectively) lag due to latency in the pathway response. Alternatively, R_1_ could increase and then decrease due to feedback mechanisms induced by sustained rapamycin treatment. We next sought to distinguish between these two possible cellular response models using the non-steady state rate approach described earlier.

To understand which mode of response cells were operating, we measured R_1_, R_2_, and R_3_ over time following rapamycin treatment. Initially, we focused on cells treated with a high concentration of rapamycin (100 nM) compared to untreated cells (DMSO). Raw autophagosomes and autolysosomes data used for rate measurements at 30 minutes are shown as an example to illustrate the procedure followed (**Fig 4A and B**). Cells were at a steady state before rapamycin treatment at 0 hours, with no changes in either autophagosome or autolysosome numbers. Following rapamycin addition, we observed an increase in autophagosome and autolysosome numbers, similar to our previous experiments. Bafilomycin A1 was then added to measure rates. This overall procedure was repeated to collect rate data from 10 minutes to 15 hours post-treatment. We first confirmed the rates of untreated cells (basal autophagy rates) were at a steady state, meaning all three rates were equal and did not vary over time (**Fig S2**).

**Figure 4.**
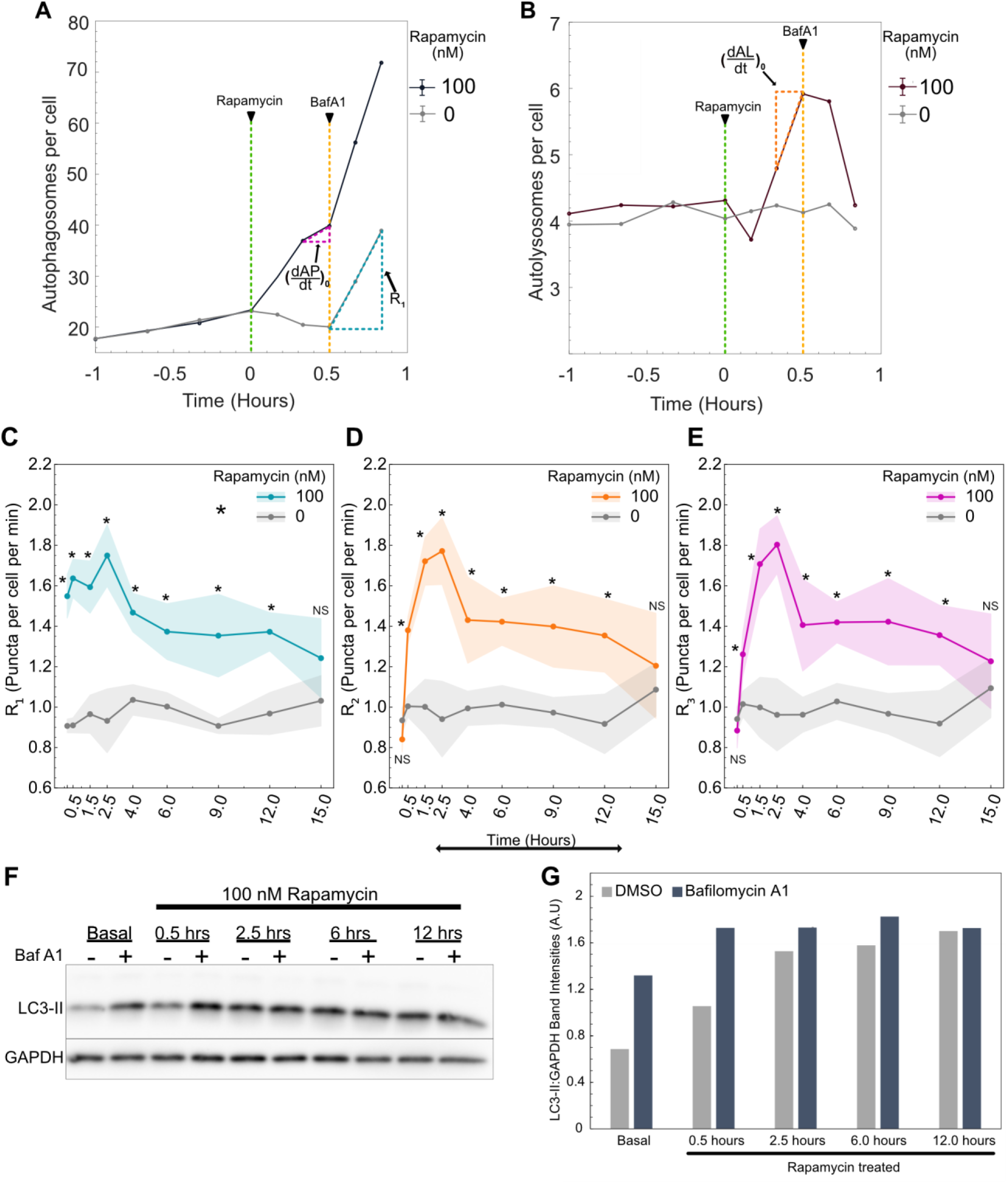
Autophagy rates change over time following rapamycin treatment. **(A)** Raw autophagosome and **(B)** raw autolysosome dynamics for rate measurement at 30 minutes post-rapamycin treatment. R_1_ was calculated using the autophagosome data 20 minutes post-bafilomycin A1 addition. 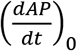 and 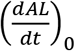 were calculated using the autophagosome and autolysosome data respectively 10 minutes before bafilomycin A1 addition. **(C)** Change in R_1_ **(D)** R_2_ and **(E)** R_3_ over time. Data points represent mean while shaded area represents ± standard deviation. Four independent replicates were performed. (*) indicates p-value < 0.05 and NS indicates not significant. P-values were calculated using an independent two-tail t-test. **(F)** LC3-II and GAPDH protein quantification using western blot. Cells were treated with 100 nM rapamycin for different time points followed by 500 nM bafilomycin A1 (Baf A1) treatment. Basal samples represent DMSO-treated cells. **(G)** Quantification of the western blot shown in Fig 4F using densitometry. LC3-II band intensity is normalized with the respective GAPDH band intensity in the same lane.

For rapamycin-treated cells, we observed a nearly immediate increase in R_1_, with significant changes in R_1_ measured as soon as 10 minutes post-treatment compared to untreated cells (**Fig 4C**). This significantly elevated rate was maintained until 12 hours post-treatment. This result is consistent with the known mechanism of rapamycin inducing autophagy upstream of phagophore elongation and thus validates the proposed non-steady state approach to characterize the effects of external perturbation on autophagy. Interestingly, R_2_ and R_3_ were slower to increase, with significantly increased rates starting at 30 minutes post-treatment (**Fig 4D and E**). Similar to R_1_, R_2_ and R_3_ maintained significantly elevated rates until 12 hours post-treatment. At 15 hours post-treatment, all rapamycin rates were statistically indistinguishable from the basal rates of untreated cells. The dynamics we observed suggest that R_2_ and R_3_ may represent rate-limiting steps initially, after which there is a general decrease in all rates.

To confirm the puncta and rate dynamics observed using the new method are consistent with the traditionally used method, we measured LC3-II protein levels using western blot. We treated cells with 100 nM rapamycin for different time points followed by the addition of 500 nM bafilomycin A1 to measure R_1_ (**Fig 4F** and **G**). Cells were treated with bafilomycin A1 for 2 hours to ensure a consistent and detectable change in the LC3-II levels. Here, the higher sensitivity of the new method is noteworthy, as it can detect changes as soon as 20 minutes after bafilomycin A1 treatment compared to 2 hours for western blot. We confirmed the increase in LC3-II levels for DMSO samples after treating with bafilomycin A1, indicating the inhibition of the fusion step. For just rapamycin treatment, we observed an increase in LC3-II levels starting at 30 minutes followed by constant maintenance of LC3-II levels until 12 hours post-treatment. This is contrary to the observed autophagosome and autolysosome puncta dynamics where there is an initial increase followed by a decrease (**Fig 3A and B**). We hypothesize that the variation between puncta dynamics measurements and western blot measurements is caused by the intrinsic nature of each measurement. LC3-II protein levels measured using western blot indicate the summation of LC3-II protein on autophagosomes and autolysosomes. Both autophagosome and autolysosome puncta remain higher than the initial state even at 15 hours, indicating higher levels of LC3-II protein at those time points. Moreover, the number of LC3-II molecules bound to each autophagosome could be dynamic and is challenging to measure. Nonetheless, the initial accumulation of LC3-II is consistent between the two methods. To validate the observed R_1_ dynamics, bafilomycin A1 was added at multiple time points after treating with rapamycin. We observed a clear increase in LC3-II accumulation at 30 minutes, while for the later time points, it only caused a modest increase. This indicates R_1_ is higher initially and declines over time. These results are consistent with measurements made using the new method where R_1_ increased over 2.5 hours post-treatment, followed by a gradual decrease.

### Latency and feedback contribute to rapamycin-driven autophagy rate dynamics

We next set out to test these temporal differences in rates by comparing rates at different time points for rapamycin-treated cells. At 10 and 30 minutes post-treatment, R_1_ was significantly greater than R_2_ and R_3_ (**Fig 5A**), consistent with the rapid increase in the autophagosome numbers until 30 minutes post-treatment. But this difference was eliminated by 1.5 hours post-treatment because of increases in R_2_ and R_3_ (**Fig 5A**), which is also consistent with the peak time of autolysosome numbers (**Fig 3B**). We focused our detailed temporal analysis on R_1_ since R_2_ and R_3_ reached the same level as R_1_ and followed the same trend from 1.5 hours onward. Interestingly, we observed a constant R_1_ until 2.5 hours post-treatment, at which point there was a gradual decrease until 15 hours (**Fig 5B**). Thus, an increase in the overall flux through the pathway in response to rapamycin is initially limited by latency in R_2_ and R_3_, but not R_1_. This is followed by a general decrease in autophagy rates after 2.5 hours. Thus, both models of regulation we initially hypothesized to exist are playing a role in the autophagy dynamics we observed.

**Figure 5.**
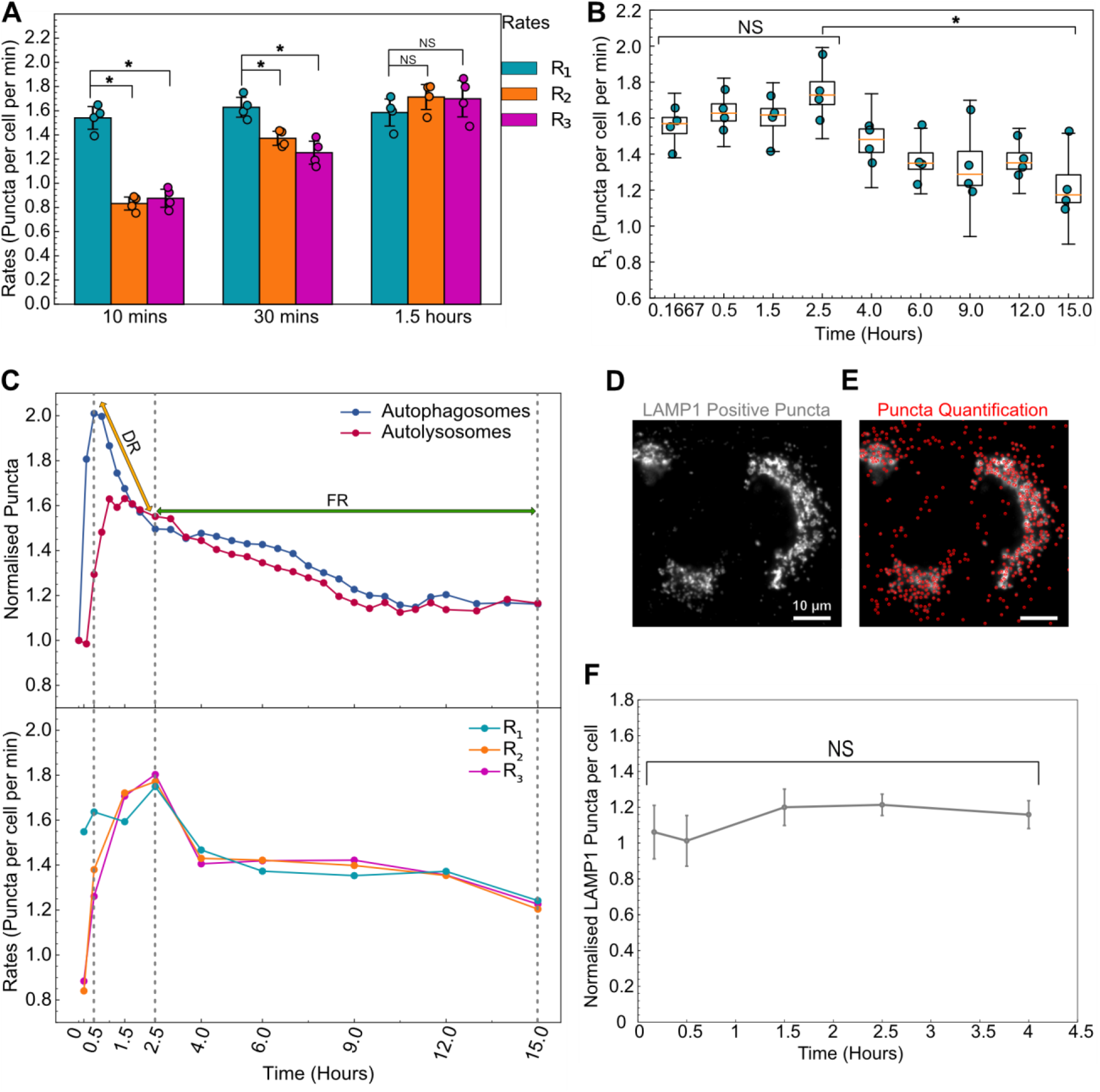
Autophagy rates indicate a hybrid model of cellular response to high concentrations of rapamycin. **(A**) Statistical comparison of autophagy rates for cells treated with 100 nM rapamycin at three different time points. **(B)** Temporal change in R_1_ for cells treated with 100 nM rapamycin. (*) indicates p-value < 0.05 and NS indicates not significant. Statistical significance for the first four points was calculated using a one-way ANOVA test. The P-value for the statistical test between 2.5 and 15 hours is calculated using paired two-tail t-test. **(C)** Normalized mean values of autophagosome (AP:AP_0_) and autolysosome numbers (AL:AL_0_) along with mean values of autophagic rates (R_1_, R_2_, R_3_) are compared to visualize the two regimes of cellular response. DR and FR represent degradative and feedback regimes, respectively. **(D)** A549 cells stained with LAMP1 antibody. **(E)** LAMP1 positive puncta detection (shown in red). Scale bar represents 10 μm. **(F)** Temporal change in the normalized LAMP1 puncta per cell (rapamycin treated to DMSO treated) after treatment with 100nM rapamycin. Error bars represent standard error for four independent replicates. NS indicates not significant, statistical significance was calculated using a one-way ANOVA test.

To illustrate this hybrid model of cellular response and regulation of autophagy rates, we juxtaposed the autophagosome and autolysosome dynamics with rate data (**Fig 5C**). We used 30 minutes as a reference point to compare the temporal changes in R_1_ since autophagosome numbers peak at 30 minutes. The immediate spike in R_1_ but lag for R_2_ and R_3_ caused the initial accumulation of autophagosomes in the first 30 minutes. After 30 minutes, the decrease in autophagosomes until 2.5 hours is caused by an increase in R_2_ and R_3_ to the same level as R_1_, leading to the degradation of accumulated autophagosomes which we name as degradative regime (DR). However, after 2.5 hours, the decrease in autophagosome numbers is a result of the decrease in R_1_ along with R_2_ and R_3_ which is labeled as feedback regime (FR). These results underscore the overall consistency of temporal rate data with the autophagosome and autolysosome dynamics.

We hypothesized that the initial lag in R_2_ and R_3_ in degradative regime is due to a lack of lysosomes to fuse with the newly formed autophagosomes. To test this hypothesis, we performed immunofluorescence staining for Lysosomal Associate Membrane Protein 1 (LAMP1) after treating cells with 100 nM rapamycin for 4 hours (**Fig 5D**). We chose 4 hours, as R_2_ and R_3_ reach R_1_ by 1.5 hours and remain the same thereafter. We quantified the LAMP1-positive puncta for basal and 100 nM rapamycin-treated cells (**Fig 5E**). Contrary to our hypothesis, there is no significant difference in the normalized puncta (rapamycin-treated to basal) between 10 minutes and 4 hours of rapamycin treatment (**Fig 5F**). This indicates that the number of lysosomes is not a limiting factor for the fusion step. Probing for the fusion governing proteins that may be rate limiting can be valuable for elucidating the fundamental mechanism involved as well as for developing new targets to inhibit the fusion step.

### Initial autophagy rates and time evolution of rates depend on rapamycin concentration

Given the concentration-dependent effects of rapamycin on autophagosome and autolysosome dynamics, we hypothesized that autophagy rates might also exhibit concentration-dependent effects. We thus measured all the rates for a range of rapamycin concentrations over 15 hours (**Fig 6A**). Mid-range concentrations of rapamycin (0.5-1.0 nM) resulted in a more gradual increase in R_1_ compared to high rapamycin concentrations (10-100 nM). To quantify rapamycin’s ability to induce autophagy, we modeled R_1_ using the Hill equation [28] (**Fig 6B**). R_1_ at 10 minutes was used to model rapamycin induction kinetics because this time point represents the effect of rapamycin on autophagy with minimal time for feedback mechanisms from the cells. In the Hill equation, (R_1_) _Basal_ represents the basal rate of autophagosome formation in the absence of any perturbation. This basal rate was estimated to be 0.90 puncta per cell per minute. V_m_ and K_m_ represent the rapamycin-induced maximal autophagy level and half-maximal rapamycin concentration, respectively. V_m_ and K_m_ are estimated to be 0.685 puncta per cell per minute and 1.1 nM. The exponent *n* represents the observed Hill coefficient and is estimated to be 1.9. This model can be used to predict rapamycin’s ability to induce R_1_ at early stages and will be useful in developing a complete temporal model. This method may be extended to various autophagy perturbations, which may relate to the mechanism of action based on the perturbed initial response and enable modeling of the response.

**Figure 6.**
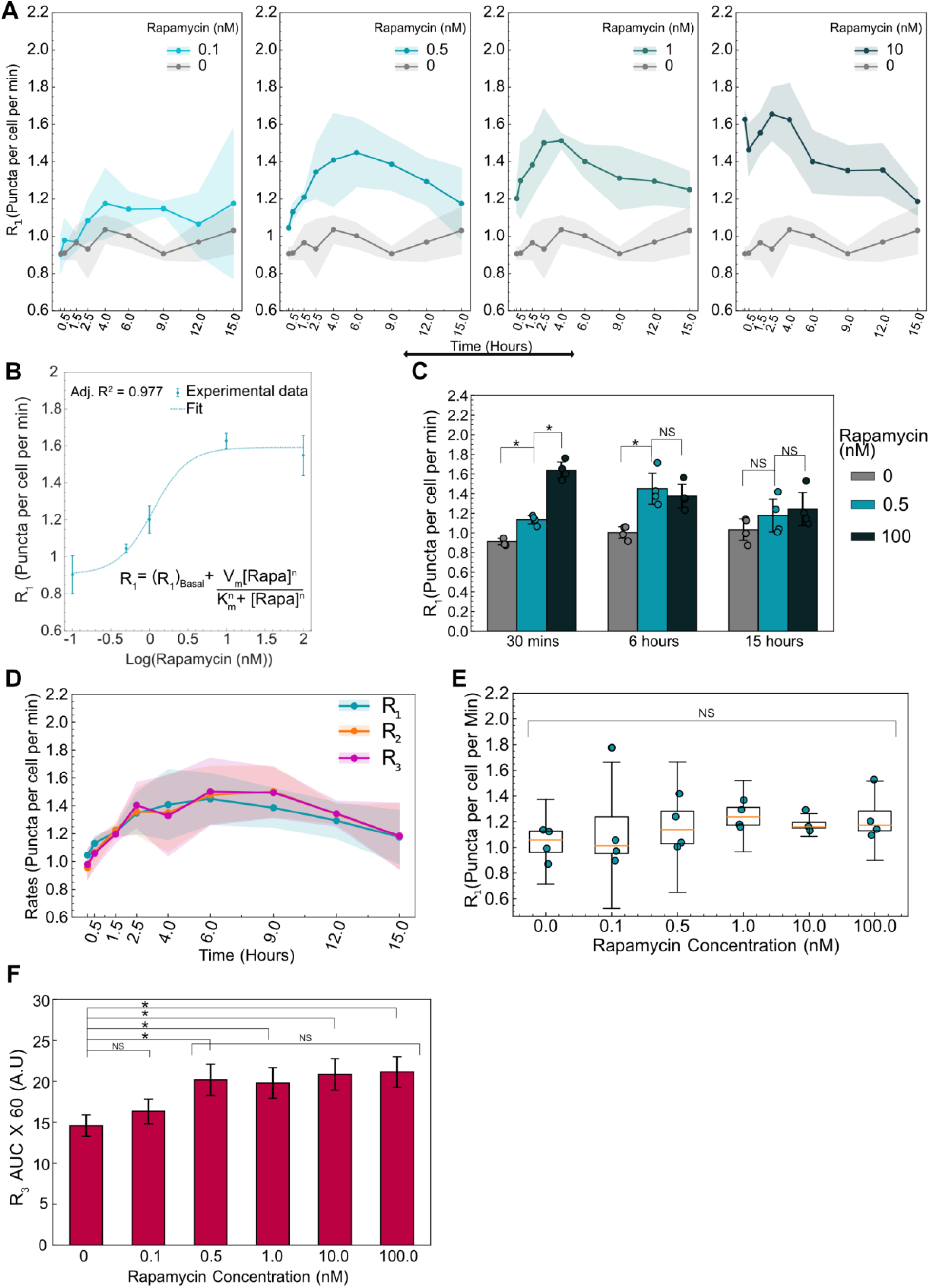
Initial and time evolution of autophagy rates depend on rapamycin concentration. **(A)** Temporal dynamics of R_1_ for different concentrations of rapamycin. Data points represent mean while shaded area represents ± standard deviation. Four independent replicates were performed. **(B)** R_1_ at 10 minutes after rapamycin addition is plotted as a function of rapamycin concentration. Individual data points represent experimental data while the dotted line represents the model fit. The model used for fitting the data along with the adjusted R^2^ value is also shown. **(C)** Statistical comparison of R_1_ for three different rapamycin concentrations at three different time points. (*) indicates p-value < 0.05 and NS indicates not significant. P-values were calculated using an independent two-tail t-test. **(D)** Temporal evolution of all autophagic rates (R_1_, R_2_, R_3_) for 0.5 nM rapamycin-treated cells. Data points represent mean while shaded area represents ± standard deviation. **(E)** R_1_ at 15 hours as a function of rapamycin concentration. NS indicates not significant. P-values were calculated using a one-way ANOVA test. **(F)** AUC for R_3_ data for different concentrations of rapamycin. Data represent mean ± standard deviation. P-values were calculated using a one-way ANOVA test followed by Tukey’s post hoc test for pairwise comparison.

In addition to concentration-dependent effects on initial R_1_, we also observed concentration-dependent effects on the time evolution of R_1_. For a mid-range concentration of rapamycin (0.5 nM), R_1_ gradually increased and reached the same level as R_1_ of higher concentrations (100 nM) over 6 hours (**Fig 6C**). This was surprising given the very low accumulation of autophagosomes and autolysosomes for 0.5 nM treatment (**Fig S1A and B**). Consequently, we explored the temporal nature of all autophagy rates (R_1_, R_2_, and R_3_) for this mid-range concentration of rapamycin. We hypothesized that the slower response time for R_1_ at mid-range concentrations of rapamycin (**Fig 6A**) might allow adequate time for R_2_ and R_3_ to adjust in sync with R_1_, even at early time points, compared to the rapid response for high concentrations of rapamycin (100 nM). Measuring R_2_ and R_3_ over time showed a similar trend, equal to R_1_ over the entire 15-hour time course, and thus resulting in low accumulation of the autophagic vesicles (**Fig 6D**). Interestingly, rates decreased at longer time points. R_1_ was indistinguishable from basal levels by 15 hours post-treatment for 0.5 nM rapamycin (**Fig 6C**), similar to the high concentration of rapamycin (**Fig 4C-E**). This led us to look at long-term impacts on R_1_ for all concentrations of rapamycin. For all concentrations tested, R_1_ was not significantly different from basal levels by 15 hours post-treatment (**Fig 6E**), suggesting adaptation of the autophagy response to long-term inhibition of mTOR complexes. Taken together, rapamycin treatment results in early responses that are concentration-dependent, with mid-range concentrations resulting in slower responses that evolve as steady state flux through the pathway. These results further signify the importance of measuring all the autophagy rates temporally to capture the complete response.

Given the temporal and concentration-dependent behavior of autophagy, we wanted to develop a simple metric for measuring the total amount of autophagy processed until steady-state conditions are reached. Assuming the average cargo captured and degraded are the same for each condition, we measured the area under the curve (AUC) for R_3_. This measure would indicate the total amount of cargo completely degraded through the autophagic pathway. For example, perturbations such as rapamycin that induce autophagic flux would have higher AUC while perturbations that inhibit autophagy initiation or clearance would have low AUC. Using the 15-hour R_3_ temporal data, we calculated the AUC for basal and all rapamycin treatments (**Fig 6F**). While 0.1 nM rapamycin is indistinguishable from the basal condition, all other rapamycin concentrations lead to significantly higher AUC, indicating higher autophagic flux. Intriguingly, all rapamycin concentrations above 0.1 nM degraded similar amounts of cargo. These results are consistent with the observed slow response for the mid-range concentration (0.5 and 1 nM) and a faster but shorter response for higher concentrations (10 and 100 nM) as discussed earlier. Henceforth, this measurement can be used as an additional metric to track the total amount of autophagy perturbed.

### Wortmannin temporarily inhibits basal and rapamycin-induced autophagosome formation

After validating the method using an autophagy inducer, we next set out to test the method using wortmannin, a commonly used inhibitor of autophagosome formation [29,30]. We used 1 μM wortmannin to test its ability to inhibit basal and rapamycin-induced autophagy. First, we measured the autophagosome and autolysosome temporal dynamics after treating with wortmannin and/or rapamycin (**Fig 7A and 7B**). We observed an immediate decrease in the autophagosome numbers for wortmannin, as well as wortmannin with rapamycin-treated conditions, while the autolysosomes numbers remained constant. Surprisingly, after 30 minutes, we observed an increase in the autophagosome numbers in cells treated with wortmannin only and a combination of wortmannin with rapamycin. Moreover, the rate of increase in autophagosome number for wortmannin with rapamycin-treated cells was faster than just wortmannin-treated cells. The autophagosomes for wortmannin with rapamycin reach initial levels by 1 hour and kept increasing to saturate at a higher level than basal (~1.8 fold) by 6 hours, followed by a slight downward trend by 15 hours. Wortmannin-only treatment takes around 3.5-4 hours for autophagosome numbers to reach the initial level and saturates at a slightly higher level (~1.3 fold). For autolysosomes, we observed a higher accumulation for wortmannin with rapamycin treatment (~1.5 fold) compared to just wortmannin (~1.1 fold), similar to the autophagosomes behavior. We used 100 nM rapamycin and basal (DMSO) as controls and their behavior remained the same as earlier experiments. From the known inhibitor mechanism of wortmannin, we hypothesize the initial drop in autophagosomes is a result of inhibition of R_1_. However, the increase after 30 minutes could either be a result of an increase in R_1_ or a much lower decrease in R_2_ and R_3_ compared to R_1_ or a combination of both. Rates for each step is needed to uncover the dynamics involved.

**Figure 7.**
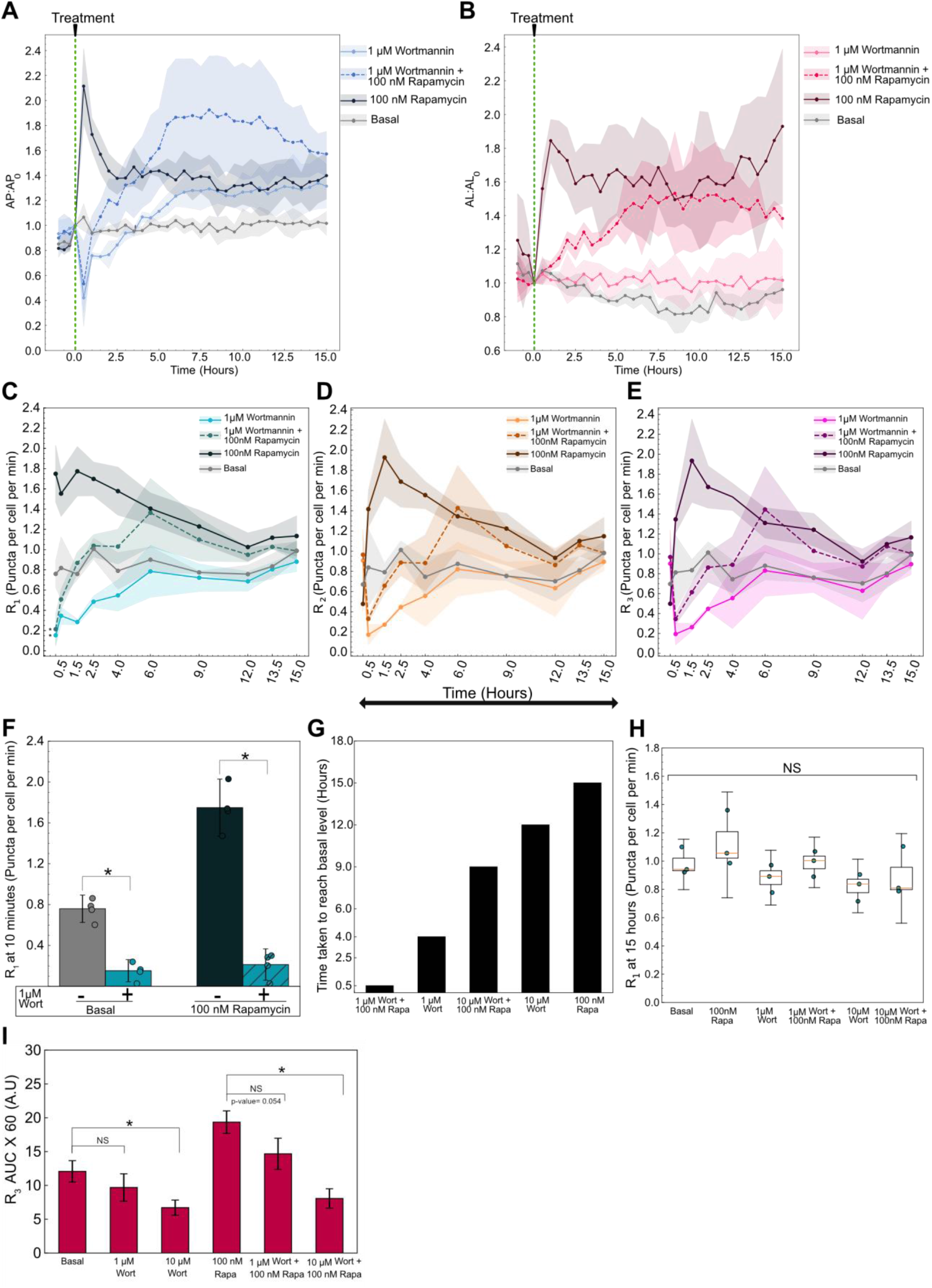
Variable autophagy recovery time from wortmannin’s inhibition. **(A)** Autophagosome and **(B)** autolysosome number dynamics after treatment. The indicated concentration of small molecule was added at 0 minutes. The number of autophagosomes and autolysosomes at 0 minutes was used as the normalization factor. Data points represent mean while shaded area represents ± standard deviation. Three independent replicates were performed. **(C)** R_1_ **(D)** R_2_ (**E)** R_3_ temporal dynamics for basal, 1 μM wortmannin (Wort) with and without 100 nM rapamycin (Rapa), and 100 nM rapamycin alone. Data points represent mean while shaded area represents ± standard deviation. Three independent replicates were performed. (*) indicates p-value < 0.05, p-values were calculated using an independent two-tail t-test. **(F)** Statistical comparison of R_1_ at 10 minutes for different treatments. (*) indicates p-value < 0.05, p-values were calculated using an independent two-tail t-test. **(G)** Time taken for R_1_ of each treatment condition to reach the basal level. (**H)** R_1_ at 15 hours plotted as a function of treatment condition. NS indicates not significant. p-values were calculated using a one-way ANOVA test. **(I)** AUC for R_3_ for different treatments. P-values were calculated using a one-way ANOVA test followed by Tukey’s post hoc test for pairwise comparison. (*) indicates p-value < 0.05 and NS indicates not significant.

We next used the non-steady-state method discussed earlier to measure the individual rates. At 10 minutes, we observed an immediate decrease in the R_1_ while R_2_ and R_3_ remained at the basal level for wortmannin-treated cells, confirming the known inhibitory mechanism of action of wortmannin (**Fig 7C-E**). Moreover, R_1_ for wortmannin with rapamycin is also significantly lower at 10 minutes compared to basal and rapamycin treatments (**Fig 7C and F**). Therefore, wortmannin initially inhibits basal as well as rapamycin-induced autophagosome formation. R_2_ and R_3_ drop by 30 minutes due to the lack of autophagosomes to degrade because of low R_1_ (**Fig 7D and E**). Interestingly, R_1_ begins to rise over time following initial inhibition by wortmannin. This could be due to either feedback from the cells and/or degradation of wortmannin over time. R_2_ and R_3_ follow a similar trend as R_1_ after 30 minutes with a slight delay (**Fig S7A and B**). This indicates that the behavior is mainly driven by R_1_ and the downstream flow of autophagosomes through the pathway is unperturbed. Rapamycin and basal rate behaviors were consistent with the previous results (**Fig 7C-E**).

We tried a higher wortmannin concentration (10 μM) to probe if it plays a role in the recovery of R_1_ (**Fig S7C-E**). We then analyzed the time taken for R_1_ to reach back to a statistically insignificant level as basal and potentially exceed it (**Fig 7H**). A treatment of 1 μM wortmannin reaches basal level by 4 hours while 10 μM wortmannin takes around 12 hours, suggesting wortmannin’s concentration is a governing factor. Moreover, we observed a faster recovery of wortmannin with rapamycin-treated cells. For 1 μM wortmannin, cells also treated with rapamycin reach basal level by 30 minutes compared to 4 hours for wortmannin alone. Similarly, for 10 μM wortmannin, the cells with rapamycin take 9 hours compared to 12 hours with wortmannin alone. The accelerated recovery of rapamycin-treated cells could be due to the additional autophagosome induction capacity of rapamycin.

We next analyzed the final steady-state rates and the total cargo degraded in terms of R_3_ AUC over 15 hours. At 15 hours, the rates of all treatment conditions were indistinguishable from each other as well as the basal condition (**Fig 7I**). This result reemphasizes the importance of temporal monitoring of autophagy, as rapamycin and wortmannin, which have opposing effects on autophagosome formation, reach the same final steady state. Finally, using the 15-hour R_3_ data, we calculated the AUC for 1 μM and 10 μM wortmannin treatment conditions (**Fig 7J**). We anticipated a decrease in the overall cargo degraded as wortmannin inhibited the initiation of autophagosome formation and thus reduced the overall flux through the pathway. We did not observe a significant decrease in the overall cargo degraded for 1 μM wortmannin treatment compared to basal but did for 10 μM wortmannin treatment. For cells treated with wortmannin along with rapamycin, 1 μM wortmannin caused a clear decrease in cargo degraded compared to the rapamycin sample even though it did not meet our statistical criteria (p-value= 0.0542). On the other hand, rapamycin with 10 μM wortmannin treatment significantly decreased the cargo degraded compared to rapamycin-induced conditions. These results are consistent with the faster recovery of autophagy under 1 μM wortmannin treatment compared to 10 μM. These measurements can be utilized to further guide the precise tuning of autophagy.

## Discussion

Quantitatively measuring all the autophagic steps remains a significant challenge and is key for developing better autophagy-based applications. This is especially critical for developing autophagy-based therapies, where dysfunction of cellular pathways is disease- and environment-specific, leading to variable response to the same treatment. Therefore, it is crucial to systematically characterize the disease state, kind of perturbation (for example, inducer or inhibitor) as well as the cellular response to gain a comprehensive understanding. Moreover, since autophagy is a dynamic process, it is pivotal to temporally monitor the process to capture the complete dynamic response until a steady state is reached. These measurements will be essential in informing the overall change in the autophagic state after a perturbation, the feedback mechanisms involved, and their timescales. For example, this information will assist in developing combinatorial therapies for effectively modulating autophagy to treat diseases with finer control and minimal side effects [31,32].

We present a method to quantify autophagy rates in live cells. Previous studies have quantified the rate of autophagosome production under steady-state conditions [12,13,18]. We expand on these studies by creating a theoretical and experimental framework to measure autophagy rates for all three steps in the autophagy pathway under non-steady state conditions. We do so by monitoring autophagosome and autolysosome numbers before and after inhibition of autophagosome-lysosome fusion. When combined with the instantaneous rate approach it enables measurement of all three autophagy rates without the requirement of them being equal.

By measuring autophagy rates under non-steady state conditions for rapamycin, we were able to validate our system using a well-characterized inducer of autophagy. We observed concentration-dependent increases in initial rates of autophagosome formation. These results were consistent with previous studies measuring autophagy flux [33,34] and rapamycin’s well-established mode of action upstream of phagophore formation. We also observed an overall return to basal autophagy rates, consistent with previous indirect observations [35]. These results are indicative of long-term feedback mechanisms at play. Importantly, our approach enables measuring initial rates with high time resolution (~10 minutes), which can uncover the direct mode of action of an autophagy perturbation, before long-term feedback mechanisms convolute measurements.

The non-steady state approach also revealed novel insights into the mechanisms regulating the cell response to rapamycin. We uncovered temporal responses to high concentrations of rapamycin that could be explained by a hybrid model of regulation of autophagy rates. Latency in the rates of autolysosome formation and degradation revealed rate-limiting steps leading to autophagosome accumulation at very early time points. Moreover, we have also shown the latency in the fusion step is not due to a limited number of lysosomes. At later time points, feedback mechanisms lowered the overall flux through the pathway. Understanding the timeline of feedback mechanisms for additional perturbations could help refine control over autophagy. On the other hand, low concentrations of rapamycin treatment led to a slower but steady response in rates. We hypothesize this behavior is due to the complex interplay between multiple feedback mechanisms of mTORC1 [36,37]. Using these measurements in conjunction with fluorescent protein activity reporters at a single cell level can elucidate the complex dynamics involved [38,39]. The temporal nature of this new approach also enabled the development of a new metric in the form of overall cargo degraded (AUC for R_3_) which can be used as an additional property to characterize the system. Using this metric, we found no significant difference in the overall cargo degraded for all concentrations above and equal to 0.5 nM rapamycin. This result underlines the dynamic nature of the pathway and the significance of this metric to fine-tune the flux through the pathway. In the future it will also be interesting to determine if autophagy-associated diseases are due to a general reduction in degradative capacity (AUC for R_3_), or defects in degradation of specific cargo.

We also measured rates for wortmannin to demonstrate the universality of this method to different types of perturbations. We observed concentration-dependent effects as well as a differential rate of recovery from wortmannin inhibition. This information can be used for modeling the system and extracting parameters such as half-maximal concentration, degradation constants, and maximal induction capacity. Additionally, this information can also be used to probe for the feedback mechanisms involved and their specific pathways. For example, if there is a mTORC1 independent feedback mechanism involved in the increase of R_1_ after wortmannin inhibition, the addition of rapamycin after recovery would lead to a higher R_1_ value and vice versa. Future work can be focused on extracting system parameters to develop a predictive model using the current data and some additional experimentation.

While our method enables novel measurements of autophagy rates, expanding its use will require improvements in autophagy-related tools. For example, our current approach uses fluorescent proteins to monitor autophagosome and autolysosome numbers and is thus limited to engineered cells. Performing similar experiments with live-cell organelle dyes could overcome this limitation. This would enable autophagy rate measurements in difficult-to-engineer cells, and open the door to measurements in patient-derived cells [40,41]. Possible applications include precision medicine for autophagy-related diseases such as cancer and neurodegeneration [42,43]. Moreover, expanding such measurements to *in vivo* systems is vital for clinical translation [44].

In conclusion, our work demonstrates quantitative measurement of rates for all three steps in the autophagy pathway under non-steady state conditions. This study revealed novel mechanisms of regulation for rapamycin induction of autophagy and differential temporal kinetics of wortmannin’s inhibition. In the future, these approaches could be applied to uncover mechanisms of action for novel autophagy-regulating compounds, develop predictive models, and characterize unique responses based on cellular genetic background. Moreover, integration with other live cell measurements would create a quantitative and holistic picture of autophagy as it connects to other cellular pathways.

## Abbreviations

MAP1LC3/LC3: microtubule-associated protein 1 light chain
LC3 ΔG: LC3 mutant lacking glycine at the carboxyl-terminus
AP: Autophagosomes
AL: Autolysosomes
t: time
R_1_: Rate of autophagosome formation
R_2_: the rate of autolysosome formation via fusion of autophagosomes with lysosomes
R_3_: Rate of autolysosome degradation
mTORC1: mammalian target of rapamycin complex 1
AUC: area under the curve
Baf A1: Bafilomycin A1
Rapa: Rapamycin
Wort: Wortmannin
GFP: green fluorescent protein
RFP: red fluorescent protein.

## Acknowledgments

We thank members of the Shah and Albeck groups for feedback on project development and the manuscript. The infrastructure used to collect data was purchased through funding by the W. M. Keck Foundation.

## Author Contributions

NSB and PSS conceived of the project, designed experiments, and wrote the manuscript. NSB performed all experiments. PSS secured funding for research.

## Materials and Methods

### Cell culture and media

A549 cells (ATCC, CCL-185) were used for all autophagy experiments. HEK 293T cells (ATCC, CRL-11268) were used for lentivirus packaging. Both cell lines were cultured in DMEM (Gibco, 11965118) supplemented with 8% fetal bovine serum (FBS [Gibco, 10438-026]). Cells were cultured in a humidified incubator at 37 °C and 5% CO_2_. For live-cell imaging, A549 reporter cells were cultured in FluoroBrite DMEM (Gibco, A1896701) supplemented with 8% FBS and 4 mM of GlutaMAX (Gibco, 35050061).

### Chemical treatments

Bafilomycin A1 (Selleck Chemicals, S1413), rapamycin (Selleck Chemicals, S1039), and wortmannin (Selleck Chemicals, S2758) were used for treating the cells. Bafilomycin A1 was added along with a final concentration of 0.2 ug/mL Hoechst 33342 trihydrochloride solution (Hoechst 33342 [Invitrogen, H3570]) to ensure proper mixing. All basal conditions were treated with DMSO (Sigma Aldrich, 472301).

### Reporter cell line construction

The FUGW-PK-hLC3 lentivirus was used to develop A549 reporter cell lines. Lentivirus was packaged in HEK 293Ts in 6-well format as previously described [45]. The harvested lentivirus media was stored at −80 °C until further use. A549 cells were plated overnight at a density of 0.1 million cells per well in a 24-well plate. The media was replaced with lentivirus-containing media. After an hour, the lentivirus media was replaced with fresh media, and the cells were scaled up upon reaching confluency. The transduced A549 cells were sorted into individual cells into a 96-well plate using Beckman Coulter “Astrios EQ”:18-Color cell sorter. Each clone population was scaled up upon reaching confluency. A single clone population was used for all the experiments to decrease noise arising from different integration sites. FUGW-PK-hLC3 ΔG reporter cell line was also developed using the same approach. FUGW-PK-hLC3 was a gift from Isei Tanida (Addgene plasmid #61440; http://n2t.net/addgene:61440). FUGW-PK-hLC3 ΔG was a gift from Isei Tanida (Addgene plasmid #61461; http://n2t.net/addgene:61461; RRID: Addgene_61461).

### Live cell microscopy

Reporter A549 cells were seeded in 96-well glass-bottom plates with #1.5 cover glass (Cellvis, P96-1.5H-N). Live cell imaging was performed using Nikon Ti2 inverted microscope with an okolab stage top incubator to maintain 37 °C and 5% CO_2_. Cells were plated at approximately 1.7 × 10^4^ cells per well for 12-18 hours prior to performing the experiment. A total of 4-5 positions were imaged in each well at the indicated time using the NIS-Elements AR software. GFP channel images were acquired at 25% LED intensity and 200 ms exposure. TRITC channel images were acquired at 30% LED intensity and 350 ms exposure settings. Images were acquired using CFI PLAN APO LAMBDA 40X CF160 Plan Apochromat Lambda 40X objective lens, N.A. 0.95, W.D. 0.17-0.25mm, F.O.V. 25mm, DIC, Correction collar 0.11-0.23 mm, Spring Loaded, and using Andor Zyla VSC-08688 camera.

### Immunofluorescence

A549s were seeded in a 96-well glass-bottom plate. Cells were treated with either DMSO or 100 nM rapamycin for indicated times. After treatment, 100% ice-cold methanol (Fisher Scientific, A412-4) was added to the cells and were incubated for 20 minutes at −20 °C. After aspirating methanol, cells were rinsed thrice with 1 X DPBS (Gibco, 21600069) solution for 5 minutes each. Following DPBS wash, cells were incubated with 5% Goat serum (Sigma Aldrich, G9023) in 1X DPBS with 0.3% Triton^™^ X-100 (Fischer Scientific, BP151-100) for an hour. After aspirating the serum solution, cells were incubated with 1:600 anti-LAMP1 primary antibody solution (LAMP1(D2D11) XP Rabbit mAb [Cell Signaling Technology, 9091], 1X DPBS, 1% Bovine Serum Albumin [BSA {Sigma Aldrich, 126609}], 0.3% Triton^™^ X-100) for overnight. After removing the primary antibody solution, cells were rinsed thrice with 1X DPBS solution. After rinsing, cells were incubated with secondary antibody solution (1:1000 Goat α-Rabbit IgG (H+L) Alexa flour 488 [Invitrogen^™^, A-11008], 1:5000 Hoechst 33342 in 1% BSA in 1X DPBS with 0.3% Triton^™^ X-100) for an hour in dark. Finally, cells were washed thrice with 1X DPBS solution for 5 minutes each before imaging. Each condition at each time point had three replicates. The average number of LAMP1 positive puncta from the triplicates was used as one biological replicate value. The experiment was repeated four times independently.

### Immunofluorescence Microscopy

After fixation, cells were imaged using Nikon Ti2 inverted microscope. LAMP1 positive puncta and nuclear staining were imaged using the green channel (GFP) and blue channel (DAPI), respectively. GFP images were acquired at 25% LED intensity and 200 ms exposure. DAPI images were acquired at 15% LED intensity and 75 ms exposure settings. Images were acquired using CFI PLAN APO LAMBDA 40X CF160 Plan Apochromat Lambda 40X objective lens, N.A. 0.95, W.D. 0.17-0.25mm, F.O.V. 25mm, DIC, Correction collar 0.11-0.23 mm, Spring Loaded, and using Andor Zyla VSC-08688 camera.

### Western blot

80, 000 cells per well were plated in a 12 well plate overnight. The next day, cells were treated with DMSO or 100 nM rapamycin for different time points. 500 nM bafilomycin A1 was added to the cells for 2 hours at different timepoints for measuring autophagic flux. At the specific time point, the media in the wells was aspirated and the cells are quickly rinsed using 1X DPBS. Cells were then lysed with RIPA buffer (150mM sodium chloride [NaCl {Fischer Scientific, S271}], 50mM Tris pH 8 [Fischer Scientific, BP152], 1% Triton X100 [Fischer Scientific, BP151-100], 0.1% sodium dodecyl sulfate [Fischer Scientific, BP166-500], 0.5% sodium deoxycholate [Sigma Aldrich, D6750]) containing protease inhibitors (Thermo Scientific^™^, A32955). The lysed cells in RIPA buffer were incubated on ice for 30 minutes. After 30 minutes, the samples were centrifuged at 12,000 RPM for 20 minutes, after which the supernatant of the samples was collected and stored at −20 °C. Sample protein content was normalized using the Pierce^™^ BCA protein assay kit (Thermo Scientific^™^, 23225). LDS (Invitrogen^™^, NP0007) and TCEP (Thermo Scientific™, 77720) in 4:1 ratio was then added to the normalized samples and were heated in a thermocycler for 10 minutes at 95 °C. The samples were then run on an SDS gel containing 4% stacking and 15% resolving gel compartments. The proteins were resolved at 115 volts for 15 minutes initially followed by 150 volts for an hour. The proteins were then transferred onto methanol-activated Amersham Hybond P 0.2 PVDF membrane (Cytiva, 10600021) at 150 volts for an hour. The membrane is then reactivated using methanol and quickly rinsed in distilled water. After reactivation, the membrane is blocked using 5% milk in TBS-T buffer (Tris-Buffered Saline pH 7.6 [TBS {20 mM Tris pH 8, 150 mM NaCl, Hydrochloric acid [Sigma Aldrich, 320331]}] with 0.1% Tween-20 [Fischer Scientific, BP337-100]) solution for an hour. The membrane slices were then incubated in their respective primary antibody diluted in 2.5% Bovine Serum Albumin (BSA [Sigma Aldrich, 12660]) in TBS-T solution for overnight with gentle agitation. The antibody dilution for each antibody is as follows, 1: 1000 anti-Phopho-S6 ribosomal protein (Ser240/244) (D68F8) Rabbit mAb (Cell Signaling Technology, 5364), 1:1000 anti-S6 Ribosomal Protein (5G10) Rabbit mAb (Cell Signaling Technology, 2217), 1:1000 anti-LC3B (D11) XP Rabbit mAb (Cell Signaling Technology, 3868), and 1: 1000 anti-GAPDH (14C10) Rabbit mAb (Cell Signaling Technology, 2118). Following primary antibody incubation, the membrane was rinsed thrice with TBS-T solution for 5 minutes each. The membrane was then incubated with 1:5000 goat Anti-rabbit IgG-HRP (SouthernBiotech, 4030-05) secondary antibody in 5% Milk TBS-T solution. The membrane was washed twice with TBS-T followed by a TBS wash. Finally, the membrane was incubated with ECL western blotting substrate (Thermo Scientific^™^, 32109) for 5 minutes before acquiring images. Amersham Imager 600 system (GE Healthcare) was used for imaging.

### Quantification and Statistical Analysis

#### Image processing for live cell imaging

NIS-Elements AR software was used for extracting autophagosomes and autolysosome puncta numbers. GFP and TRITC channel images were processed and analyzed using the General analysis job functionality in the NIS-Elements AR software. Both GFP and TRITC channel images were background corrected using the rolling ball correction method with a radius of 1.95 *μ*m. Following background correction, Spot Detection functionality was used for thresholding and detecting puncta in both channels. GFP channel images were used for estimating autophagosome puncta numbers as the green signal is only detected in autophagosomes. Bright-clustered detection method in the Spot Detection tool was used for detecting circular areas in the GFP channel with a typical spot diameter of 0.8 *μ*m and a minimum contrast value of 5. The contrast value acts as a thresholding parameter to only detect puncta whose difference between mean intensity inside and mean intensity outside the spot is higher than the contrast value provided. Similarly, for the TRITC channel, 0.8 *μ*m was used as the typical spot diameter and 7.5 was used as the contrast value. A higher contrast value was used for detecting puncta in the TRITC channel because of the lower signal-to-noise ratio and thus to minimize false-positive puncta. However, the puncta from the TRITC channel includes both autophagosomes as well as autolysosomes count as both have a red signal. Therefore, to extract autolysosome-only count, we compared the colocalization of puncta in GFP and TRITC channels using the AND binary operation. The number of colocalized puncta (representing autophagosomes) were then subtracted from the total TRITC puncta, thus providing the autolysosome count. All the other parameters in the spot detection tool were left as default. The puncta detection accuracy was confirmed through manual inspection of multiple images under various conditions (untreated, rapamycin and bafilomycin A1 treatment). Post analysis, the autolysosome and autophagosome count for each image were exported as an excel sheet.

Cellpose was used for counting cells in each image [46]. The ND2 files were converted to RGB tif files and the GFP channel images were used for segmenting and extracting the cell count. Cellpose was implemented in python 3.7 using a custom-written script and 120 was used as the diameter input for segmenting individual cells. The segmentation accuracy was confirmed by manual inspection as well as by comparing with Hoechst-based nucleus count.

After extracting the cell, autophagosome and autolysosome count from each position imaged in a well. The total number of autophagosomes and autolysosomes in all the positions imaged were added and was divided by the total number of cells providing a population level autophagosome and autolysosome count per cell. This analysis was done using a custom-written script in MATLAB.

#### Image processing for immunofluorescence microscopy

LAMP1 positive puncta in fixed samples were estimated using spot detection tool in NIS-Elements AR software. A radius of 0.8 μm and a contrast value of 10 was used for detecting LAMP1-positive puncta. Similarly, the number of cells was estimated using the spot detection tool on the nuclear stain with a typical diameter of 15 μm and a contrast value of 1.5 as parameters.

#### Data fitting and area under the curve estimation

Curve fitting toolbox in MATLAB was used to fit the data. Custom equations were provided for fitting the data and the Nonlinear least squares method was used for the fit. Trapz function in MATLAB was used for calculating the area under the curve and a custom-written script was used for propagating the error.

#### Statistical analysis

At least 150-200 cells were imaged for all experiments. A minimum of three experimental replicates was performed for all the quantitative experiments. Independent t-test, paired t-test, and one-way ANOVA were used as indicated for comparing statistical significance for various experiments. All statistical tests were performed in python 3.8 using the SciPy package. ANOVA along with Tukey post hoc test for AUC calculations was performed using a webpage (https://statpages.info/anova1sm.html). Box and whisker plots indicate the median value as an orange line, interquartile range (IQR) as a box, and range [*Q*_1_ – 1.5 * *IQR*, *Q*_3_ + 1.5 * *IQR*] as whiskers.

## Supplemental Information

**Figure S1:**
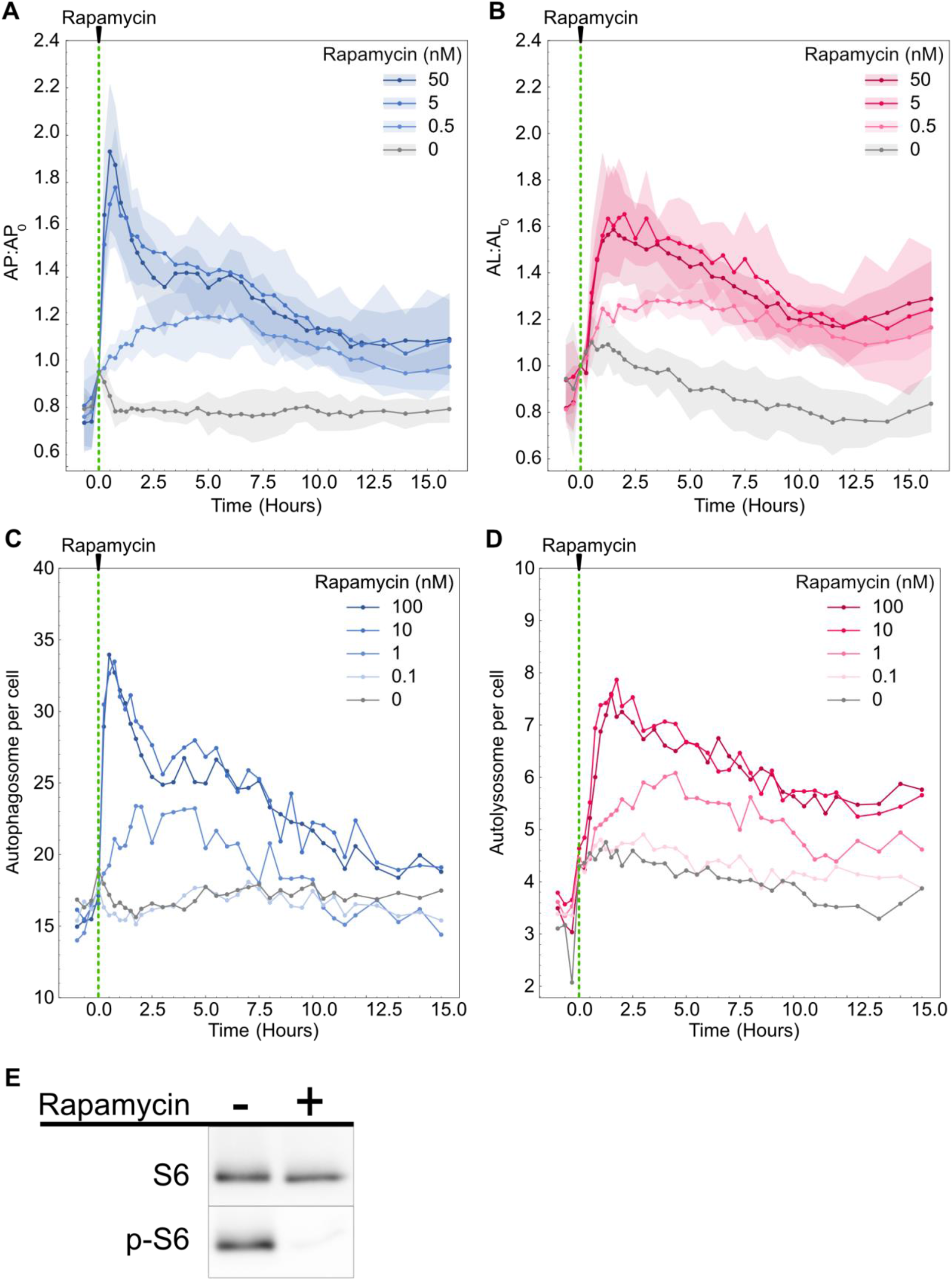
Autophagosome and autolysosome dynamics for additional rapamycin concentration. Dynamics of **(A)** Autophagosome and **(B)** autolysosome number after rapamycin treatment. The indicated concentration of rapamycin was added at 0 minutes. The number of autophagosomes and autolysosomes at 0 minutes was used as the normalization factor. Data points represent mean while shaded area represents standard deviation. Four independent replicates were performed. Raw **(C)** autophagosome and **(D)** autolysosomes dynamics of one individual replicate after rapamycin treatment (**E**) Change in phosphorylation of phospho-S6 ribosomal protein (Ser 240/244) (p-S6) after treatment with 100 nM rapamycin for 2.5 hours. S6 ribosomal protein (S6) is used as a control.

**Figure S2:**
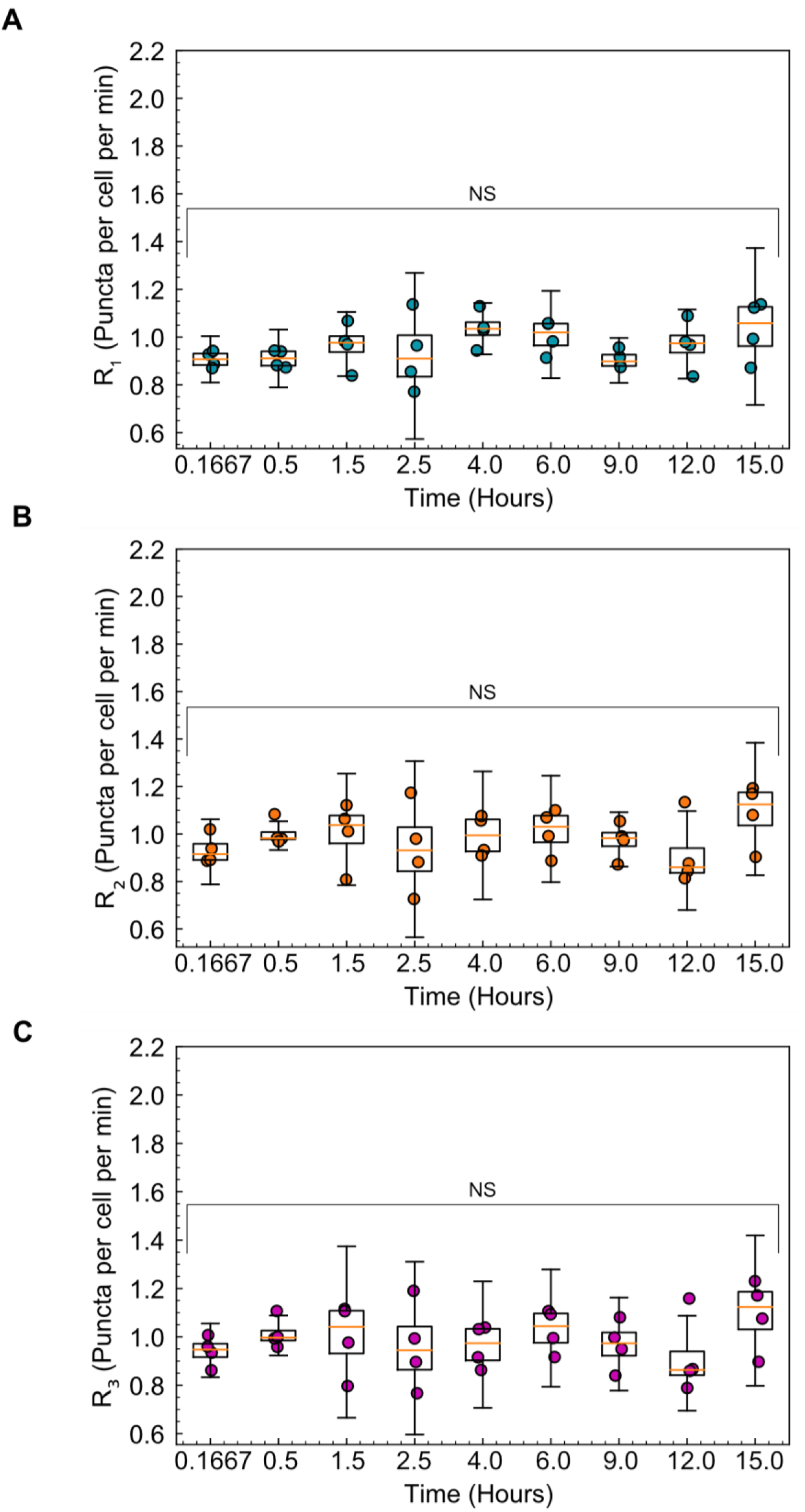
Basal autophagy rates remain constant over time. **(A)** Rate of autophagosome formation (R_1_). **(B)** Rate of autolysosome formation (R_2_). **(C)** Rate of autolysosome degradation (R_3_). NS indicates not significant. P-values were calculated using a one-way ANOVA test.

**Figure S3:**
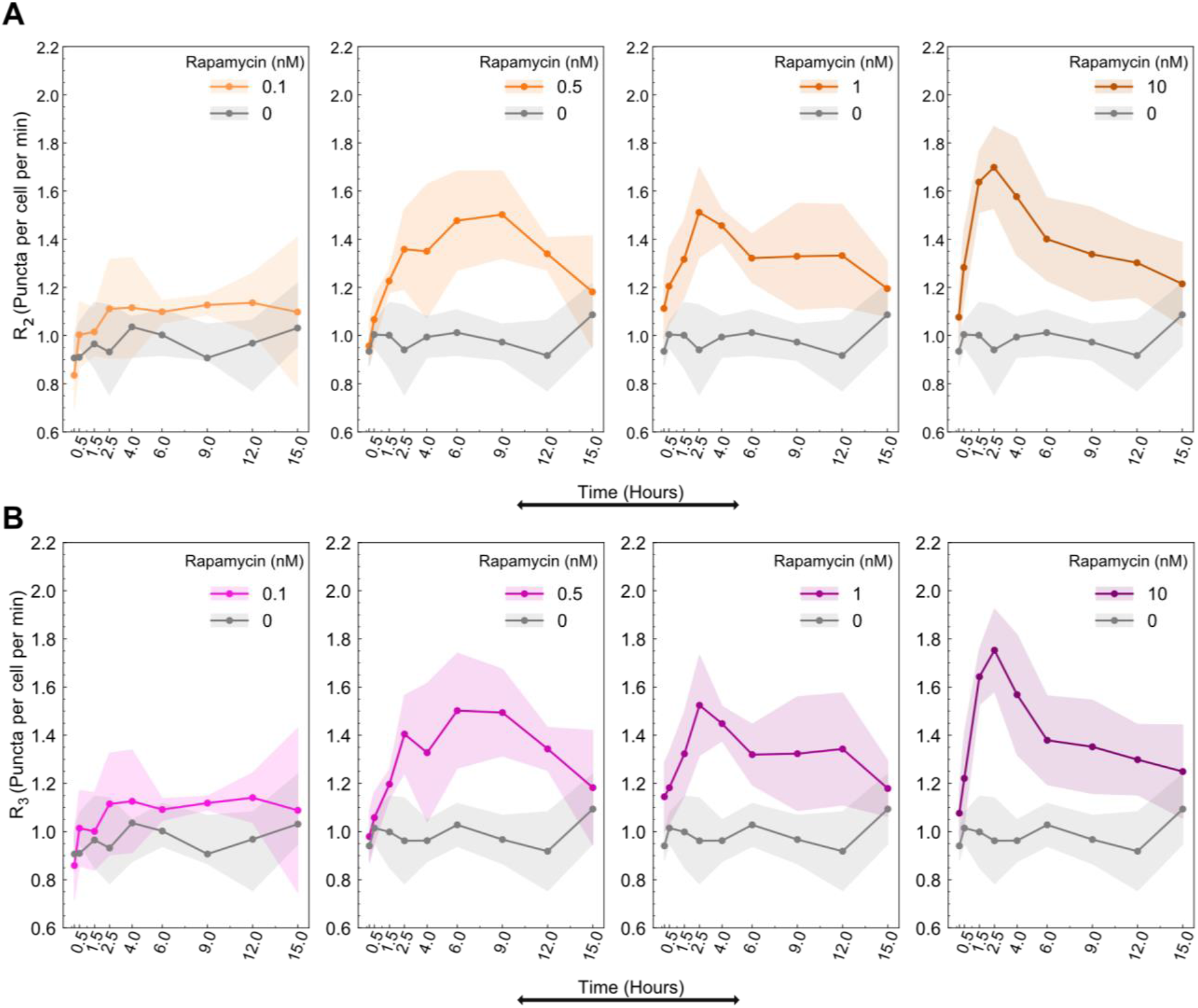
Rapamycin concentration modulates autolysosomes formation and degradation rates. **(A)** Temporal dynamics of rate of autolysosome formation (R_2_) and **(B)** Autolysosome degradation (R_3_) for different concentrations of rapamycin. Data points represent the mean, while the shaded area represents ± standard deviation. Four independent replicates were performed.

**Figure S4:**
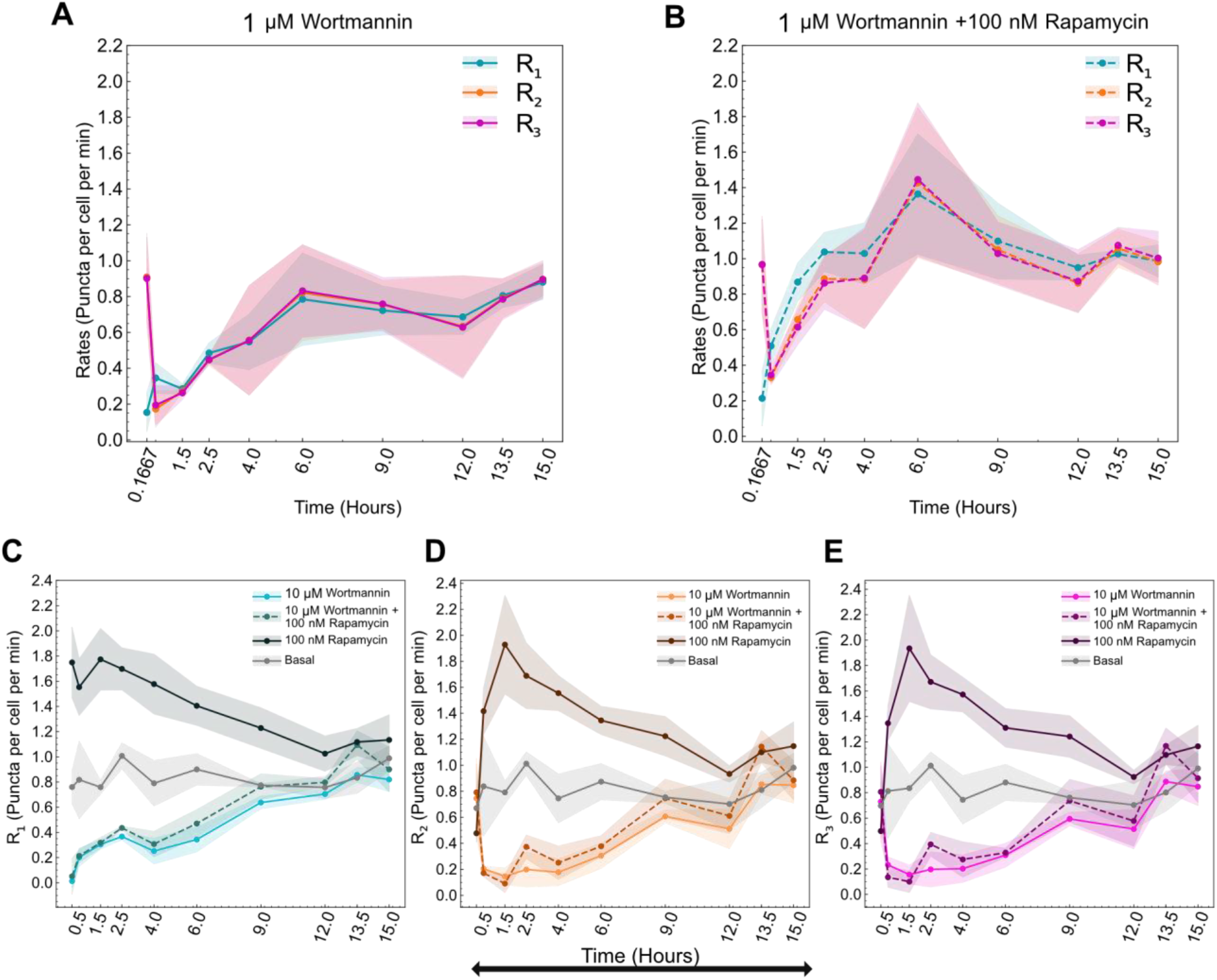
Autophagy temporal dynamics during wortmannin treatment. Temporal evolution of all autophagic rates (R_1_, R_2_, R_3_) for **(A)** 1 μM Wortmannin only and **(B)** 1 μM wortmannin with 100 nM Rapamycin. Temporal dynamics of **(C)** R_1_ **(D)** R_2_ (**E)** R_3_ for basal, 10 μM wortmannin with and without 100 nM rapamycin, and 100 nM rapamycin alone. Data points represent the mean, while the shaded area represents ± standard deviation. Three independent replicates were performed.

